# Functional and effective connectivity between dorsolateral prefrontal and subgenual anterior cingulate cortex depends on the timing of transcranial magnetic stimulation relative to the phase of prefrontal alpha EEG

**DOI:** 10.1101/2022.02.14.480466

**Authors:** Spiro P. Pantazatos, James R. Mclntosh, Golbarg T. Saber, Xiaoxiao Sun, Jayce Doose, Josef Faller, Yida Lin, Joshua B. Teves, Aidan Blankenship, Sarah Huffman, Robin I. Goldman, Mark S. George, Paul Sajda, Truman R. Brown

## Abstract

**BACKGROUND:** The communication through coherence model posits that brain rhythms are synchronized across different frequency bands and that effective connectivity strength between interacting regions depends on their phase relation. Evidence to support the model comes mostly from electrophysiological recordings in animals while evidence from human data is limited.

**METHODS:** Here, an fMRI-EEG-TMS (fET) instrument capable of acquiring simultaneous fMRI and EEG during noninvasive single pulse TMS applied to dorsolateral prefrontal cortex (DLPFC) was used to test whether prefrontal EEG alpha phase moderates TMS-evoked top-down influences on subgenual, rostral and dorsal anterior cingulate cortex (ACC). Results in healthy volunteers (n=11) were compared to those from patients with major depressive disorder (MDD) (n=17) collected as part of a ongoing clinical trial investigation.

**RESULTS:** In both groups, TMS-evoked functional connectivity between DLPFC and subgenual ACC (sgACC) depended on the EEG alpha phase. TMS-evoked DLPFC to sgACC effective connectivity (EC) was moderated by EEG alpha phase in healthy volunteers, but not in the MDD patients. Top-down EC was inhibitory for TMS onsets during the upward slope of the alpha wave relative to TMS timed to the downward slope of the alpha wave. Prefrontal EEG alpha phase dependent effects on TMS-evoked fMRI BOLD activation of the rostral anterior cingulate cortex were detected in the MDD patient group, but not in the healthy volunteer group.

**DISCUSSION:** Results demonstrate that TMS-evoked top-down influences vary as a function of the prefrontal alpha rhythm, and suggest clinical applications whereby TMS is synchronized to the brain’s internal rhythms in order to more efficiently engage deep therapeutic targets.

## 1 INTRODUCTION

In the communication through coherence model, cognition relies on patterns of neural synchronization that change dynamically with stimulation and behavioral context (1, 2). Brain rhythms are synchronized across different frequency bands and are highly structured across areas, layers and their corresponding projections (1). The model proposes that bottom-up-directed gamma band influences are controlled by top-down-directed alpha-beta band influences, and that rhythmic modulation of postsynaptic excitability and coherence between pre- and postsynaptic neuronal groups are the basis for strong effective connectivity (1). Moreover, the effective connectivity strength between interacting regions may depend on their phase relation, as suggested by recordings in visual cortex of awake cats and monkeys (3).

Whether the phase of endogeneous EEG rhythms mediate top-down influences in the human brain, particularly in response to neurostimulation, is an active area of investigation. If so, it could have a large impact in both basic and clinical neuroscience. EEG, fMRI, and transcranial magnetic stimulation (TMS) have been used together in a pair-wise fashion to investigate brain networks, and the feasibility of concurrent fMRI-EEG-TMS (fET) with offline analysis of results has been previously demonstrated (4). We have created a system of simultaneous fET, where the timing of TMS delivery relative to the EEG signal can be determined. This enables assessment of EEG phase-dependent TMS effects on brain activity. Importantly, it would allow us to test whether TMS that is synchronized to the brain’s internal cortical rhythms has differential effects on deeper corticolimbic brain structures involved in emotion processing and regulation. If so, this could facilitate clinical applications. For example, TMS could be timed to optimize therapeutic engagement of the anterior cingulate cortex (ACC) in patients with major depressive disorder (MDD).

Previous research using simultaneous TMS-EEG has demonstrated that motor excitation following TMS pulses varies with EEG phase and amplitude (5, 6, 7, 8). Using simultaneous EEG-fMRI, our group has previously shown that BOLD activity in decision related networks were modulated by the prestimulus EEG alpha phase during an auditory oddball task (9). Based on these previous findings, we hypothesized that single pulse TMS stimulation synchronized with respect to alpha phase would modulate ACC functional and effectivity connectivity with the stimulation site as well as ACC activation.

Here, fET was used to determine whether TMS delivery to DLPFC during different phase points in the cycle of alpha oscillations modulates functional and effective connectivity with ACC. Whether alpha phase of TMS onset modulated ACC BOLD activity was also examined. Our fET instrument can acquire simultaneous fET and remove fMRI artifacts automatically in real-time (10). Offline analyses were used to recover the exact phase of alpha during each single pulse TMS onset via acausal signal processing. Being able to observe changes in the level of connectivity and activity in the ACC following a TMS pulse that is timed to EEG cortical rhythm may lead to optimizations in TMS timing that could have better clinical efficacy than current, non-synchronized TMS approaches. We analyzed three subregions of ACC (dorsal ACC, or dACC; rostral ACC, or rACC, and subgenual ACC, or sgACC) based on their previously established functional or structural connectivity with DLPFC and relevance to TMS treatment of depression and related symptomatology (11, 12, 13).

## 2 RESULTS

### 2.1 Functional connectivity between DLPFC and sgACC depends on EEG alpha phase

#### ROI analysis

For generalized Psychophysiological Interactions (PPI) analysis (see methods and materials section 5.4.3), TMS onsets were grouped according to four phase bins (bin 1: −*π* to 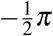, bin 2: 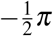 to 0, bin 3: 0 to 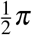, and bin 4: 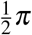 to *π*). In both datasets, alpha phase at TMS onset moderated DLPFC-sgACC functional connectivity (**HV**: F_3,40_ = 3.07, *p* = 0.038 uncorrected; **MDD**: F_3,64_ = 3.2, *p* = 0.029 uncorrected; Fisher’s combined *p* = 0.009), but not DLPFC-dACC or DLPFC-rACC functional connectivity (see Figure 1, top row). The DLPFC-sgACC result was confirmed with non-parametric p-values using a permutation based approach (**HV**: *p* = 0.02; **MDD**: *p* = 0.02; Fisher’s combined *p* = 0.004).

**Figure 1.**
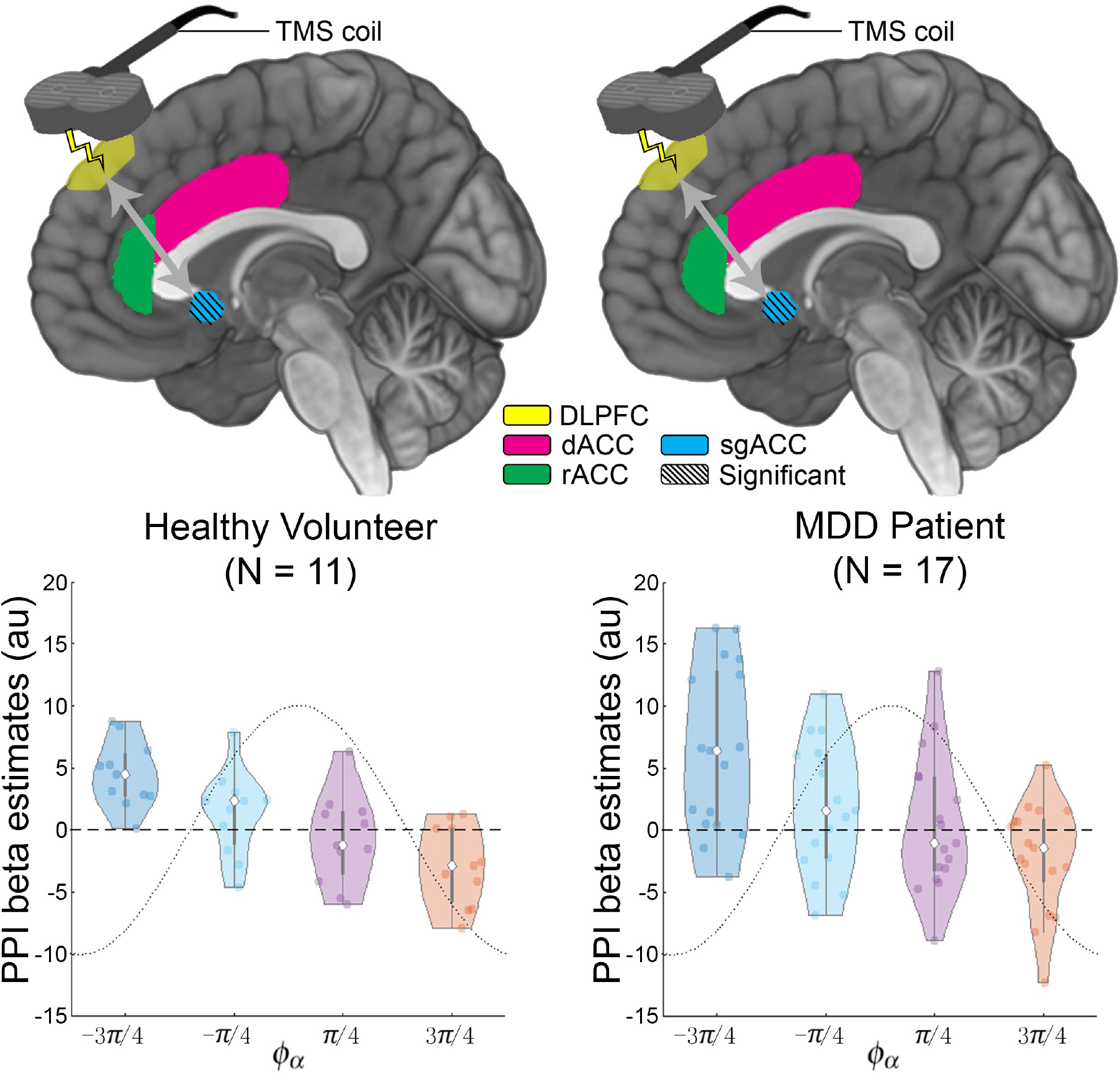
Alpha phase at TMS stimulation moderates functional connectivity between DLPFC (EEGF3 stimulation site) and subgenual anterior cingulate cortex (sgACC). Generalized psychophysiological interactions analysis (PPI) was used to estimate TMS-evoked DLPFC-sgACC functional connectivity within each of four alpha phase bins. The voxel-wise analysis was restricted to a search space within a subgenual ROI generated using a 10mm sphere centered at MNI=[6 16 −10] and excluding white matter as in (12). A group level omnibus F-test on the beta weight parameter estimates (PE) identified a cluster in the primary dataset (left: *p* < 0.05 corrected). The findings replicated in the second independent cohort (*p* < 0.05 corrected, right). The bottom row shows violin plots of PEs averaged within clusters (contiguous voxels surviving *p* < 0.05 uncorrected).

#### Small Volume Correction

In addition to conventional ROI analysis (where signal within each mask is averaged together), a voxel-wise analysis within each ACC mask was applied using the mask for small volume correction. Consistent with the ROI analysis results, alpha phase bin at TMS onset moderated functional connectivity between the DLPFC TMS target site and sgACC in the **HV** dataset (voxel-wise FWE corrected *p* = 0.068, cluster-extent corrected *p* < 0.05). A cluster within the sgACC ROI search region was defined using an F-test thresholded at uncorrected *p* < 0.05, then used to visualize and plot the beta parameter estimates for each of the four phase bins across subjects. The functional connectivity between DLPFC and sgACC was highest (most positive) when the TMS onset occurred during the rising phase of the alpha oscillation in bins 1 and 2 (−3/4*π* and −1/4*π*), and the lowest (negative) during the falling phase in bins 3 and 4 (centered at 1/4*π* and 3/4*π*, respectively, see Figure 1, bottom left). This finding replicated in the **MDD** dataset (voxel-wise FWE corrected *p* = 0.037, cluster-extent corrected *p* < 0.05, see Figure 1, bottom right). The greatest pairwise difference, consistent across both datasets, was between phase bin 1 vs. bin 4: DLPFC−sgACC functional connectivity was highest in bin 1 and lowest in phase bin 4 (**MDD**: t_64_ = 4.4, FWE voxel-wise corrected *p* = 0.004, cluster extent corrected *p* < 0.05).

### 2.2 Top-down DLPFC-sgACC effective connectivity depends on EEG alpha phase

Functional connectivity results indicate there is a statistical association between fMRI BOLD activity in ACC and DLPFC that depends on alpha phase, but they do not enable inference about the direction of information flow between the two areas. To infer causal influences between DLPFC and ACC in response to TMS, as a function of alpha phase, a simple dynamic causal model (DCM) of DLPFC and sgACC was estimated for each subject. The DCM modeled experimental modulation (TMS) of both extrinsic and intrinsic connections during each of four phase bins (see section 2.1). A form of Bayesian Model Comparison and Reduction (14) was applied to compare the full model to ‘reduced’ versions of the model along three factors: TMS modulation of top-down vs bottom up extrinsic connectivity, TMS modulation of DLPFC vs sgACC intrinsic connectivity, and TMS as driving input to DLPFC vs sgACC (see Figure S.1A in supplementary information, SI). The full DCM model is represented by the 1^*st*^ model along each factor shown in Figure S.1A in SI. This created a search space of 64 unique models. The average ‘baseline’ effective connectivity (EC), or bidirectional connectivity representing the mean across all TMS phase conditions or during baseline in non-centered models, *A* matrix in the DCM (15), was included in all models (see methods section 5.4.4).

Model evidence was summed across all models belonging in each level, or ‘family’ of models, for each factor (Figure S.1B in SI). In the **HV** dataset, results suggest a model in which TMS modulates intrinsic connectivity of both DLPFC and sgACC as well as both bottom-up and top-down connectivity between the two regions, while TMS does not directly drive activity in either region. In the **HV** dataset, the average top-down effective connectivity (EC) was excitatory (i.e. the *A* matrix parameter was positive with significant posterior probability *P_p_* > 0.95), and the bottom-up EC was inhibitory (negative *A* matrix parameter, *P_p_* > 0.95), while in the **MDD** dataset, both the top-down and bottom-up average EC were excitatory (Figure 2A). In the **HV** dataset, the top-down EC was modulated by TMS alpha phase (uncorrected *p* < 0.001), while trend-level evidence suggested intrinsic sgACC connectivity was modulated by TMS alpha base in both datasets (Figure 2A). A violin plot of the top-down EC in the **HV** dataset shows higher (more excitatory) connectivity when TMS was delivered during alpha phase bins 3 and 4 (bin centered at *π*/4 and 3*π*/4, respectively, Figure 2B, left), while a plot of intrinsic (self) sgACC connection in the **MDD** dataset shows TMS is excitatory (less inhibitory) when delivered during phase bin 1 (−3*π*/4) relative to the other phase bins (Figure 2B, right). Note the self-connections are unitless log scaling parameters that scale (multiply up or down) the default connection strength of −0.5Hz, so the more negative the self-connection parameter, the less inhibited the region.

**Figure 2.**
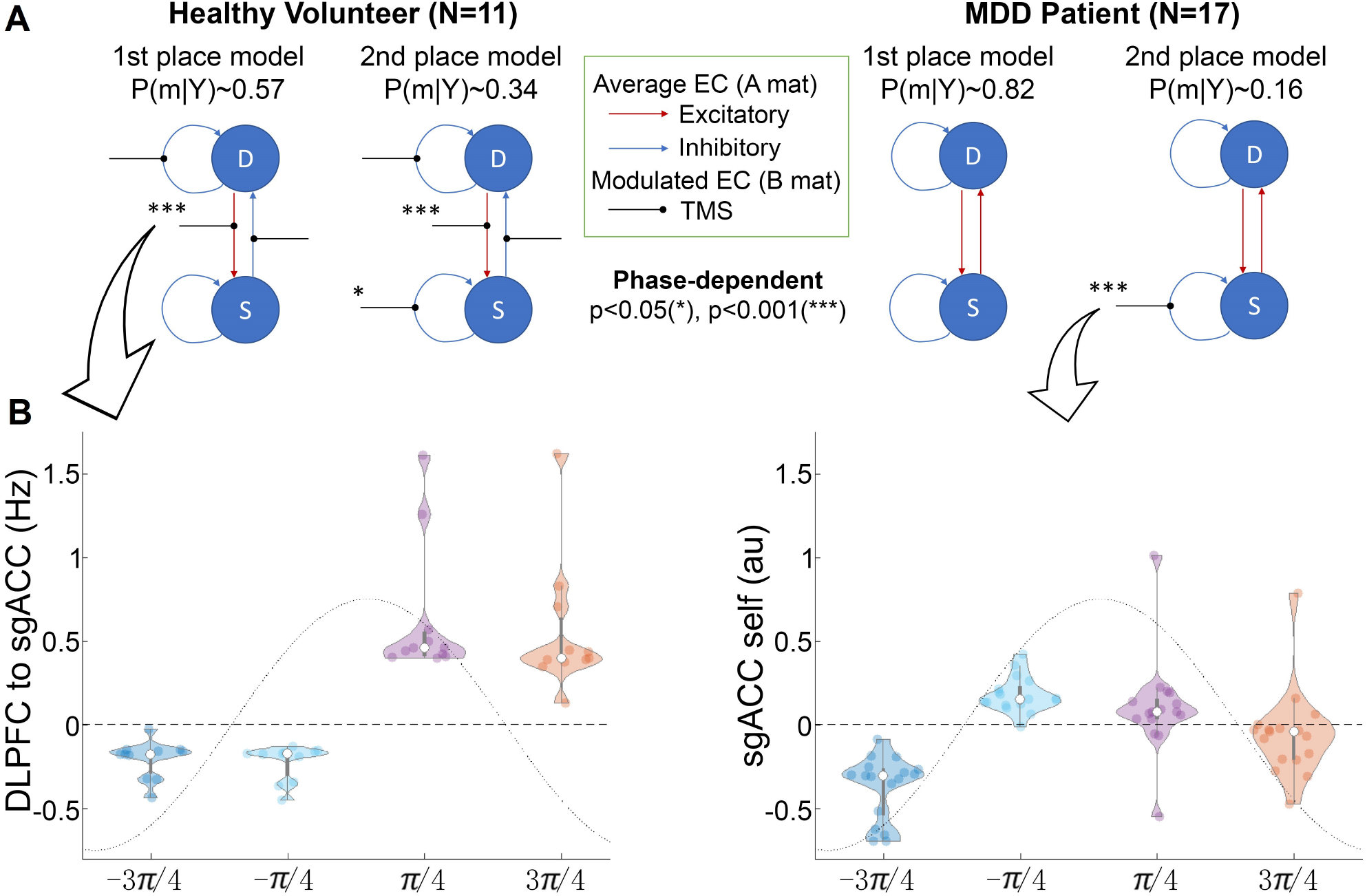
Dynamic causal modeling suggests TMS-evoked top-down DLPFC-sgACC effective connectivity (EC) depends on EEG alpha phase. **A)** The top 2 winning network structures from Bayesian Model Comparison (see methods) are shown in each dataset. EC strengths that were modulated by alpha phase at TMS onset are shown with asterisks (* indicates *p* < 0.05 uncorrected, *** indicates *p* < 0.001 uncorrected). Results suggest prefrontal alpha phase moderates TMS-evoked DLPFC-sgACC top-down EC in HV, but not in MDD. B) Violin plots in HV show top-down EC are inhibitory during the rising phase of alpha, and excitatory during the falling phase. In MDD, the DCM results suggest alpha phase at TMS onset moderates intrinsic (self) sgACC connectivity, with the lowest parameter values (excitatory effects) during the initial rising phase of the alpha wave. See Figure S.1 in SI for network model space and model comparison results.

The results presented in Figure 2B reflect models with mean centered input matrix *U* (see (15), where the inputs are TMS onsets with zero duration during the four conditions (phase bins). According to (15) this can increase the model evidence, by enabling the connectivity parameters to stay nearer to their prior expectation (of zero) during model fitting. It also means the parameters in matrix *A* represent the average effective connectivity across experimental conditions. Our primary analysis mean centered the inputs since we were mainly interested in testing for differences in connectivity strengths across the four TMS phase bins.

We also examined the effects of not mean centering the inputs as we were interested in estimating the effects of TMS modulation on connectivity, in general, relative to the implicit baseline (i.e. periods in between TMS onsets). This also allowed us to further explore sgACC-DLFPC feedback loop dynamics by estimating effective connectivity during the baseline periods (in between TMS stimulation). Results from these models are presented in Figure S.1 of SI. When the input matrix *U* was not mean centered, the extrinsic (top-down and bottom-up) connectivity strengths no longer contributed significantly to the model fit. In these models, TMS modulated intrinsic (self) connections of DLPFC and sgACC, while sgACC intrinsic connectivity was modulated by phase bin (see Figure S.2 in SI). The most consistent finding across both sets of models (mean-centered and non-mean centered input matrices *U*) was that models whereby TMS modulated intrinsic sgACC connectivity had the best model fits, and that the intrinsic sgACC connectivity strengths showed a phase-dependent effect.

### 2.3 fMRI BOLD activation of the rACC depends on EEG alpha phase at TMS onset in MDD patients

When applying conventional ROI analysis of TMS-evoked ACC fMRI BOLD activation, no effect of phase bin was detected in any of the ROIs in either dataset (corrected *p* > 0.2, see Supplementary Results S.1 and S.6 for TMS-evoked BOLD signal averaged over subjects and runs for each phase bin). We therefore applied Generalized Additive Mixed Model (GAMM, see section 5.4.5) which allows the use of smooth nonlinear functions to detect relationships between alpha phase and ACC fMRI BOLD activation. The approach is more sensitive in detecting such non-linear relationships since it provides a smooth fitting across alpha phase (i.e., avoids binning of the alpha phases). Table 1 shows the main effect of phase and phase by region interactions. In Table 1, the approximate significance of smooth terms based on alpha phase (*ϕ_α_*) is listed. In the **HV** dataset (N=11), we did not detect a significant main effect for phase-dependent BOLD activation across the 3 ACC subregions (*f* (*ϕ_α_*), *p* = 0.246) and there was no difference between the three regions (*p* > 0.05) from the GAMM result. In the (**MDD**) dataset (N=17), the phase-dependent BOLD activation was significant in rACC (*f* (*ϕ_α_*):rostral, *p* = 0.017 uncorrected). Applying Fisher’s method to combine p-values across results from both datasets suggested an overall phase-dependent BOLD activation effect within rACC (*p* = 0.040). The results with respect to brain regions in the **MDD** group are summarized in Figure 3 (left panel), while Figure 3 (right panel) plots the corresponding normalized BOLD signal (BOLD_*nor*_) differences between −*π* and *π* in rACC.

**Figure 3.**
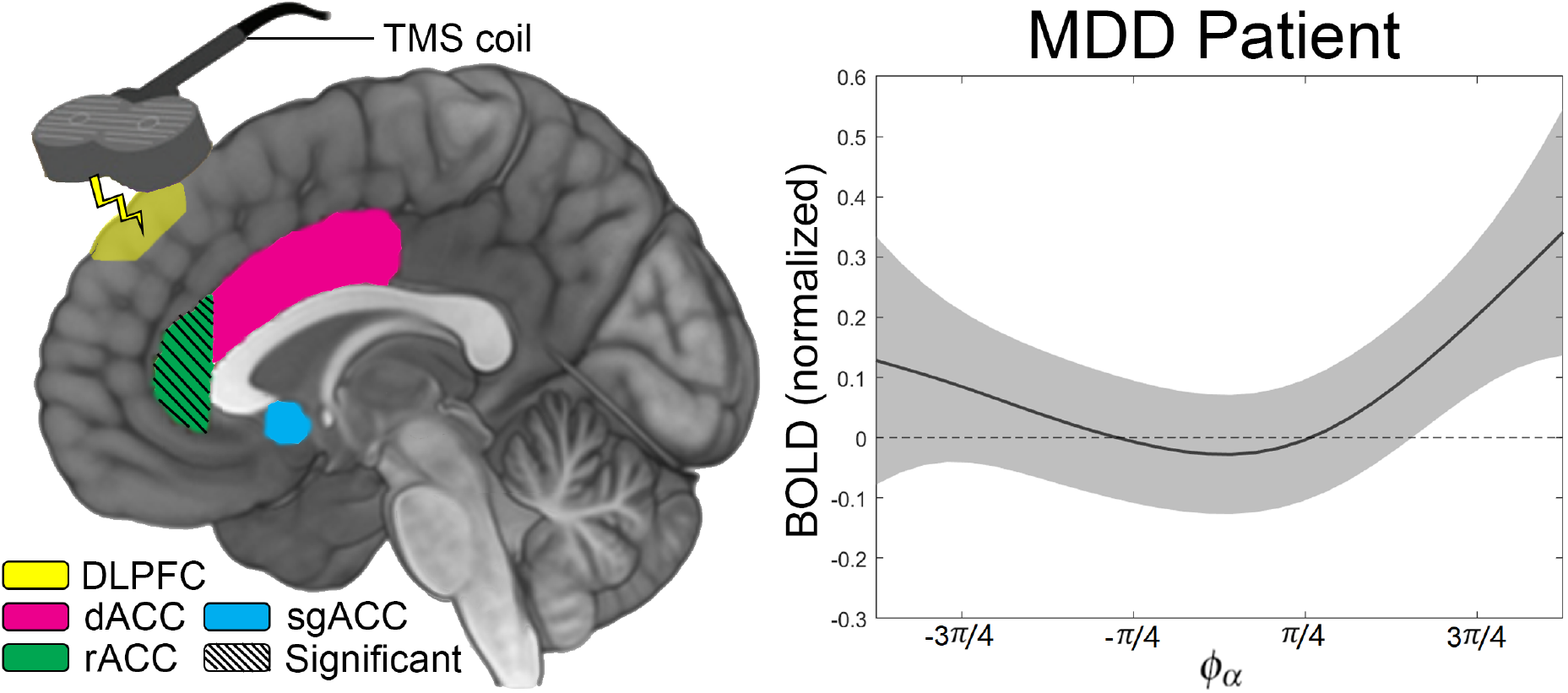
TMS-evoked BOLD activation in rACC is correlated with alpha phase at TMS onset in the **MDD** dataset. The phase-dependent BOLD activation is significant (diagonal stripes) in rACC (left panel). The right panel plots the BOLD activation (curves fitted from the GAMM models) against alpha phase. The results show that in the **MDD** dataset, the TMS-evoked BOLD response in the rACC (i.e. the normalized BOLD activation *BOLD_nor_*, see methods section 5.4.5) is related to the corresponding alpha phase *ϕ_α_* at which the TMS pulse is triggered (see Table 1, random effects on the subject level are not plotted).

**Table 1:**
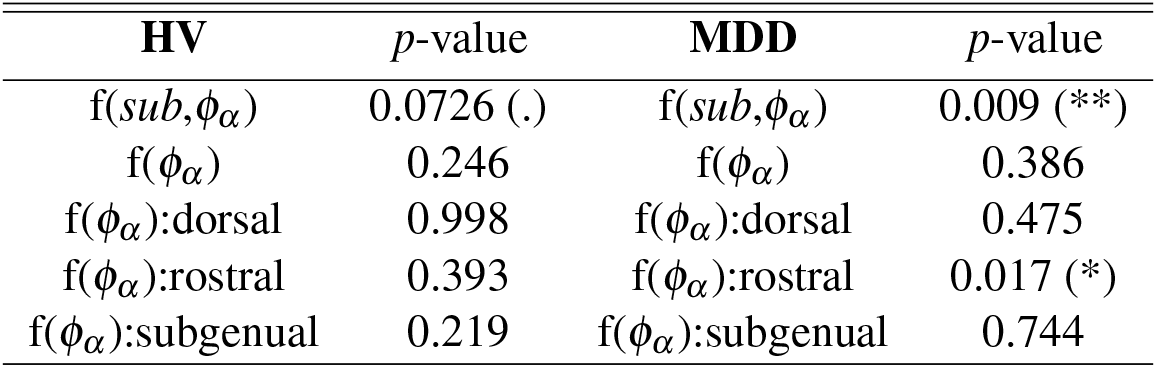
GAMM results in **HV** and **MDD** datasets and approximate significance of smooth terms. (**) indicates significance at the 99% confidence level; (*) indicates significance at the 95% confidence level; (.) indicates significance at the 90% confidence level.

| **HV** | *p*-value | **MDD** | *p*-value |
| --- | --- | --- | --- |
| $f(sub, \phi_\alpha)$ | 0.0726 (.) | $f(sub, \phi_\alpha)$ | 0.009 (**) |
| $f(\phi_\alpha)$ | 0.246 | $f(\phi_\alpha)$ | 0.386 |
| $f(\phi_\alpha):dorsal$ | 0.998 | $f(\phi_\alpha):dorsal$ | 0.475 |
| $f(\phi_\alpha):rostral$ | 0.393 | $f(\phi_\alpha):rostral$ | 0.017 (*) |
| $f(\phi_\alpha):subgenual$ | 0.219 | $f(\phi_\alpha):subgenual$ | 0.744 |

## 3 DISCUSSION

In this study, we used our recently developed integrated fET instrument capable of acquiring simultaneous EEG-fMRI while delivering noninvasive single pulse TMS, and found that TMS applied to DLPFC modulates the region’s connectivity with ACC in a way that depends on prefrontal EEG alpha phase. In both the **HV** and **MDD** datasets, phase dependent functional and effectivity connectivity with the DLPFC stimulation site was seen in sgACC. The most consistent phase-dependent effect, observed in both datasets, was DLPFC-sgACC functional connectivity following single pulse TMS. The EEGF3 DLPFC-sgACC TMS-evoked functional connectivity straddles zero and is overall weakly negative, consistent with resting state studies that found EEGF3 DLPFC-sgACC functional connectivity is the least (weakly) negative (*r* ≈ −0.05) compared to other TMS stimulation sites in the DLPFC (i.e. see Figure 1 in (12)). With the GAMM model, we also observed phase dependent activation in the **MDD** group, such that TMS-evoked BOLD responses in the rACC were lowest when TMS pulses were synchronized nearer to the peak of the frontal alpha oscillation.

An effective connectivity analysis suggests that in **HV** there is an excitatory/inhibitory feedback loop between DLPFC and sgACC and that TMS has a phase dependent effect on the top-down connection. The effective connectivity analysis in the **MDD** dataset suggests: (1) the DLPFC-sgACC feedback loop is purely positive (excitatory connections) and reflects hyperactivation of sgACC/ACC; (2) the phase dependent effect of TMS is not seen in the top-down connection (as in the **HV** group), though there are phase-dependent effects on sgACC intrinsic connectivity. The observed functional and effective connectivity effects could be mediated via indirect structural connections to sgACC or via direct structural connections between DLPFC and sgACC. Tract tracing studies in primates have shown sparse structural connections between left posterior DLPFC and sgACC (16)).

A study of single unit recordings in V1 of awake monkeys estimated spike probability as a function of gamma phase LFP, and found highest (lowest) spike probabilities during the peak (trough) of the gamma wave (1, 17). Our GAMM results followed a similar pattern, whereby fMRI BOLD signal was lowest for TMS onsets timed nearest to the peak of the alpha wave. However, functional and effective connectivity did not line up with the peaks and troughs of the EEG alpha wave. Rather, these measures lined up with the slope of the EEG alpha wave, whereby functional connectivity was highest (positive) for TMS onsets on the upward slope of the alpha wave, and lowest (negative) for TMS onsets during the downward slope of the alpha wave. The DCM analysis (in **HV**) suggested TMS-evoked top-down DLPFC-sgACC EC was negative (inhibitory) for TMS onsets on the upward slope of the alpha wave, and positive (excitatory) for TMS delivered during the downward slope of the alpha wave. The results are consistent with a previous study that found neocortical activity converged to a preferred phase of 45 degrees in response to exogenous or endogenous electrical fields (18). The findings suggest that the DLPFC is “primed” to propagate signals to sgACC when TMS is delivered during the rising phase of prefrontal alpha.

### 3.1 Clinical application in depression

The effects observed here have potentially important implications for clinical applications involving TMS. Depression involves distributed cortico-limbic networks, with anterior cingulate cortex (ACC) playing an important role in the etiology and treatment of the disorder (19, 20). TMS targeting left DLPFC is currently used for treatment of medication-resistant depression (21, 22, 23, 24), yielding remission rates of 28 to 53% (25). Understanding the mechanisms of rTMS and increasing the efficacy of rTMS is an important area of research.

The anti-depressant effects of TMS are likely mediated by connectivity between DLFPC and ACC, and the modulation of ACC activity (26, 12, 27). Previous studies have examined TMS efficacy based on spatial location of stimulation within DLPFC and find that anterolateral DLPFC regions lead to better outcomes perhaps because they are more negatively functionally connected with sgACC (28). Additionally, several groups have reported that alpha-band oscillations are associated with both activation and inhibition across and within different functional networks (29, 30, 31). Therefore, by controlling or targeting the phase of alpha rhythms, we could potentially activate or inhibit brain excitability within certain networks that are phase-dependent.

Our study finds that TMS-evoked brain excitability or functional connectivity can be indexed by phase. This suggests that deep targets may be more efficiently engaged by TMS via optimizing TMS delivery based on these indeces (e.g., alpha phase) of excitability or functional connectivity, which in turn could improve TMS treatment efficacy. In other words, TMS treatment could be improved by synchronizing the timing of TMS stimulation with the brain’s internal EEG rhythms in way that maximizes excitability or functional connectivity with deep therapeutic targets (32, 10, 33).

## 4 CONCLUSION

Using simultaneous fMRI-EEG-TMS (fET) we find that the timing of TMS relative to prefrontal alpha phase results in differential sgACC functional and effective connectivity with the DLPFC stimulation site. TMS applied during the rising phase of the alpha wave has an inhibitory effect on sgACC, while TMS during the falling phase has an excitatory effect. In addition, we find evidence for EEG alpha dependent BOLD activation in the rACC, at least in the **MDD** group. The findings demonstrate that prefrontal alpha wave activity modulates top-down influences from executive brain regions to anterior cingulate. Moreover, they suggest important clinical applications whereby TMS that is timed to internal brain rhythms could lead to more effective rTMS and improve clinical outcomes. Future work will examine whether rTMS that is synchronized to alpha wave activity can improve clinical outcomes.

## 5 METHODS AND MATERIALS (SUPPLEMENTAL ONLINE

### 5.1 Subjects

The primary dataset consisted of eleven healthy volunteer participants (**HV**: 5 male, 6 female; average age (years) 30.5 ± 8.8). Written informed consent was obtained from all subjects prior to the experiment and our experimental protocol was approved by the Institutional Review Board of the Medical University of South Carolina. All subjects were screened for history of epilepsy or medications that may lower seizure threshold.

The second dataset consisted of seventeen treatment resistant depressed patients (**MDD**: 5 male, 12 female; average age (years) 48.2 ± 13.5). The patients were recruited as part of a clinical trial. Written informed consent was obtained from all subjects prior to the experiment and the experimental protocol was approved by the Institutional Review Board of the Medical University of South Carolina. All subjects had Hamilton scores greater than or equal to 20 at enrollment (average 30.2, range 20 to 39). Further details on subject inclusion and exclusion criteria can be found in the SI.

### 5.2 fMRI-EEG-TMS (fET) acquisition

The study employed a simultaneous 3-way fET acquisition system developed by our group and previously described in (10). The instrument included a custom 43 channel MR compatible bipolar EEG system (Innovative Technologies, CA, USA), a Siemens 3T Prisma MRI Scanner using a custom 12 channel head coil (Rapid MR International), and a MagStim Rapid2 TMS neurostimulator with a modified MRI-compatible TMS coil. Prior to the scanning session, motor threshold was measured on the subject’s left motor cortex adjusting the TMS output voltages until an involuntary thumb twitch was observed. The TMS coil was configured to a subject-specific intensity of 100 to 120 % of the motor threshold.

Functional imaging was performed with a custom 12-channel head coil (Rapid MR International, LLC, Columbus, OH, USA) and a multi-echo multiband pulse sequence (CMRR, University of Minnesota). In the **HV** dataset, a high resolution structural T1 image was acquired for registration as well as a field map for distortion correction using the custom 12 channel head coil. In the **MDD** dataset, the high resolution structural T1 image was collected using a 32 channel Siemens head coil in a separate session prior to a second session for the 3-way fET acquisition.

Sequence parameters for the **HV** dataset were as follows: voxel size = 3.2 × 3.2 × 3.2 mm, slices number = 36, MB acceleration factor = 2, repetition time (TR) = 1600 ms, TE1 = 11 ms, TE2 = 32.16 ms, TE3 = 53.32 ms. Inter-pulse interval was pseudorandomized and uniformly distributed from four to six TRs. Four to six runs were collected for each subject yielding 184 (4 × 46) to 276 (6 × 46) TMS pulses per session. Sequence parameters in the **MDD** dataset were as follows: multiband acceleration factor = 2, TE1 = 11.20 ms, TE2 = 32.36 ms, and TE3 = 53.52 ms. Whole-brain fMRI was acquired in 38 slices, voxel size 3.2 × 3.2 × 3.2 mm, TR = 1.75 s, flip angle 58 degrees, with a 200ms TR gap. Reverse phase encode sequences were acquired to unwarp the images.

In both datasets, an MR compatible bipolar EEG cap with 36 electrodes was placed on the subject’s head. Individual impedances were reduced with electrode gel to below 20kΩ. The location of the DLPFC was measured and marked (under the F3 electrode) for TMS coil placement in the MRI scanner. The TMS coil was run through a filter into the MRI room and placed over subjects’ DLPFC outside the EEG cap. The EEG cap was connected to a MRI-compatible amplifier that sampled at 488 Hz with a gain of 400. A custom 12 channel head coil was used to acquire both the anatomical and BOLD scans. EEG was simultaneously acquired at 488 Hz sampling rate. All system clocks were synchronized with the 10 MHz scanner clock. The scanner trigger initiated the EEG acquisition for each run. TMS pulse firing and timing were automatically controlled via a custom Eprime (Psychology Software Tools) program synchronized with the scanner trigger. Each TMS pulse was fired at the beginning of a 200ms gap at the end of each TR. The gap allowed for the magnetic effects of the TMS pulse to not affect subsequent image acquisition. Subjects were instructed to keep their eyes open and look at a fixation cross during all runs.

### 5.3 EEG data processing

EEG data was converted to EEGLAB formatted files from the raw amplifier binary files (ITEK) before preprocessing. TR markers and TMS pulse markers were marked accordingly. The ITEK amplifier being used is severely perturbed by each TMS pulse displaying a decaying amplitude low frequency oscillation of approximately 1 Hz. In order to remove this artifact in each channel, the time series between two TMS pulses is extracted and then fitted with multiple functions: an offset function, a decaying exponential function, a sinusoidal function, and a decaying exponential modulated by a sinusoidal waveform. Oscillation removal was done by subtracting the best fitting model of those functions. Then, gradient artifact removal was done to remove the artifacts introduced by fMRI acquisition. A standard template subtraction approach (FMRIB tools) (34, 35) was used followed by removal of residual principal components, however template construction was performed on an augmented signal to reduce the impact of the artifacts induced by TMS. The data used to construct the template was augmented by replacing individual TRs that contained remaining low frequency oscillation content, TMS pulses, or amplitudes exceeding 2.5mV with a previous uncontaminated TR.

The ballistocardiogram (BCG) artifact is likely to corrupt phase estimation of the alpha rhythm. Typical BCG artifact removal methods construct a template aligned to the detection QRS events (34, 36). In this study, BCG removal was not applied because the corruption of phase induced by the BCG is unlikely to be systematic in relation to the timing of TMS pulses, while the mislabeling of QRS events as TMS pulses with standard algorithms can lead to a systematic phase change in the vicinity of the TMS pulses.

Bipolar recordings were converted to a unipolar configuration by application of the shortest path function (37), with the right mastoid set as reference. The right mastoid was chosen to match the application described in previous work (10).

Estimating the phase of the alpha oscillation *ϕ_α_* at the trigger time of a TMS pulse is challenging because with conventional non-causal methods, the TMS artifact inevitably corrupts the phase estimate. To resolve this, alpha phase (*ϕ_α_*) was estimated with a machine learning approach that produces causal estimates of the instantaneous phase from the raw signal (38). In brief, a participant specific FIR filter (0.5s, 488Hz) was constructed with cutoff frequencies centered around the peak amplitude frequency as detected with the FOOOF toolbox (39) applied to a 5 minute resting state recording taken prior to the main experiment. The peak frequency was estimated in a range from 7.5 to 12 Hz in order to avoid major components of fMRI artifact frequencies. The cutoffs were set to be −2Hz to +2Hz, or twice the central alpha peak standard deviation when modeled as a Gaussian. The average of channels FP1, F3 and F7 was used as the source of the alpha signal. Sections of this signal that did not overlap with the TMS events were acausally filtered and Hilbert transformed. The real and imaginary components were then used as targets for a regularized (L2, *λ* = 2 × 10^−4^) linear regression with the Toeplitz matrix of the backwards difference of the EEG channel average as input (see McIntosh and Sajda (38)). The two learned filters of 244 weights were then applied to causally predict the phase at the TMS event times with EEG data preceding the TMS event (see Figure S.3 in SI for the EEG preprocessing flow chart).

### 5.4 MRI data preprocessing and analysis

#### 5.4.1 Regions-of-interest (ROI) definition

Three regions (dorsal, rostral and subgenual ACC) were defined and used for ROI functional connectivity and activation analysis. Dorsal ACC (dACC) and rostral ACC (rACC) anatomical masks were generated from freesurfer in AFNI for the left and right dorsal (caudal) and rostral (ventral) ACC. The subgenual ACC (sgACC) ROI was generated using a 10mm sphere centered at MNI=[6 16 −10] and excluding white matter as previously described (12). This location corresponds to the MNI coordinate that is most responsive to interventions for depression averaged across multiple studies (12), and is also anticorrelated with various TMS DLPFC target sites at rest (28).

#### 5.4.2 Data preprocessing

The T2-weighted fMRI data were preprocessed using SPM12. For each of six sessions, the runs from three multiecho sequences were combined (averaged) into a single 4D image. The functional images were then realigned (motion corrected), slice-time corrected, spatially transformed to a standardized brain (Montreal Neurologic Institute), resampled to 2 × 2 × 2mm resolution, then smoothed with a 5-mm full-width half maximum Gaussian kernel.

#### 5.4.3 Functional connectivity

##### 1^*st*^ level fMRI analysis

A conventional SPM analysis that modeled phase dependent BOLD activation was conducted to facilitate subsequent functional connectivity (PPI) analyses. Four regressors-of-interest were created by convolving the onset of each TMS delivery pulse with the canonical HRF with duration of 0 seconds, corresponding to alpha phases binned into four equally spaced bins covering a full period beginning 0.15s prior to TMS pulse. The four bins were centered on the following values: −3*π*/4 (bin1∈ [−*π,* −1/2*π*)), −*π*/4 (bin2∈ [−1/2*π,* 0)), *π*/4 (bin3∈ [0, 1/2*π*)), and 3*π*/4 (bin4∈ [1/2*π, π*]). Nuisance covariates included 6 motion parameters, cerebrospinal fluid (CSF) and white matter mean signal, as well as spike confound regressors from fsl motion outliers (default parameters and “DVARS”) to adjust for volumes corrupted by large motion (40). Note in “DVARS” the D refers to the temporal derivative of the timecourses, and VARS refers to RMS variance over voxels (40). Peri-stimulus time histograms (PSTH) were also plotted using finite impulse response (FIR) models with time bin of one TR in Marsbar v0.44 following (41, 42) and averaged across subjects.

##### Psychophysiological Interactions (PPI) analysis

Generalized PPI analysis (43, 44) was used to test whether alpha phase moderates functional connectivity between the DLPFC TMS target site and the three ROIs defined above (dACC, rACC and sgACC). The seed region (EEGF3 target site in DLPFC, hereafter referred to as DLPFC) consisted of a 10mm sphere centered at MNI=[-37 26 49]. Four regressors-of-interest (representing connectivity with DLPFC during the four alpha phase bins) were constructed by taking the product of the DLPFC time course (physiological variable) and the vector of the TMS onset times in each of the four phase bins (i.e., 1×bin1 + 0×bin2 + 0×bin3 + 0×bin4 for phase bin1, 0×bin1 + 1×bin2 + 0×bin3 + 0×bin4 for phase bin2, etc.). Note that with generalized PPI, the interaction is assessed at the contrast level through statistical inference on the differences in beta weights reflecting the regression slopes estimated for each phase bin, see (43)). The model also included covariates representing the main effects of the physiological variable (DLPFC time course) and psychological variable (TMS onsets convolved with the canonical HRF) to adjust for “baseline” DLPFC-ACC connectivity and to remove any covariation between regions due to activation in response to the TMS. Additional nuisance covariates included CSF and white matter mean signal, and spike confound regressors from fsl motion outliers 1 (default parameters and dvars) to adjust for volumes corrupted by large motion (40). The resulting PPI beta maps for the four regressors of interest were subsequently passed to 2^*nd*^ level random effects analysis.

##### 2^*nd*^ level analysis

One-way ANOVA models (dependent factor: phase bin with four levels) were conducted separately on the beta images generated from the 1st level PPI analysis. An omnibus F-test was used to test for any difference between the four phase bins in both datasets. In addition, the resulting effects of interest plot for the significant cluster within the search sphere of the **HV** dataset was used to generate a directional hypothesis for a T-test in the **MDD** group which was treated as a replication dataset.

##### ROI analyses and correction for multiple comparisons

Two types of ROI analyses were conducted. The first was a conventional ROI analysis using Marsbar v0.44 (http://marsbar.sourceforge.net) in which signal (i.e., voxel-wise beta estimates from 1^*st*^ level analysis) within each ROI was first averaged together before model estimation. Five ROIs were included in this analysis (sgACC and left and right rACC and dACC). The uncorrected p-values obtained from each of the 5 ROIs/models were submitted to a ‘meta-analysis’ across the primary and replication datasets using a Fisher’s combined p-value test.

In the second type of analysis, three ROIs (sgACC, bilateral rACC and dACC) were used as a masks for small volume correction (SVC) in which models were estimated across all voxels within the mask. The SVC analysis was used in conjunction to the traditional ROI analysis in order to increase sensitivity for signal arising from subregions within the ROI masks that could have been averaged away when using the first ROI analysis type. For the SVC analyses, results were corrected for multiple comparisons within each mask using voxel-wise Family-Wise Error rate (FWE) correction at *p* < 0.05.

To increase sensitivity to weaker, spatially distributed signal, cluster-wise correction was applied using AFNI 3dClustSim (compiled December 11, 2018) with the -acf option (input parameters estimated using residuals from the 2^*nd*^ level SPM model), and a cluster determining threshold of *p* < 0.01. For each ROI, this yielded a minimum cluster size above which clusters were considered significant at *p* < 0.05 corrected when using 3^*rd*^ nearest-neighbor (corners touching) clustering. This approach was based on recent recommendations to reduce false-positive rates when using cluster-extent correction (45, 46) and includes a more accurate estimate of the noise smoothness values using a mixed model (Guassian plus a monoexponential) (47). For sgACC minimum cluster size was *k* = 21, dACC minimum cluster size was *k* = 20, and rACC minimum cluster size was *k* = 22.

We further confirmed conventional parametric p-values obtained from the traditional ROI Marsbar analysis by computing “non-parametric” p-values using permutation. For this, SPM .mat design files were randomly permuted across subjects 50 times to create a null distribution of F-values for the main effect of alpha phase. This shuffling scheme effectively kept onsets of TMS delivery the same, while shuffling the alpha bin labels. For the **HV** dataset, the shuffling was conducting in 8 of 11 subjects since 3 of the subjects had less then 6 total sessions. Non-parametric p-values obtained across the two datasets were then combined to compute a “non-parametric” Fisher’s combined p-value.

#### 5.4.4 Effective connectivity analysis

Group effective connectivity analyses were applied using Dynamic Causal Modeling (DCM) and two level Parametric Empirical Bayes (PEB) using DCM v12.5 toolbox for SPM12 following (15, 48). Under this framework, a single ‘full’ DCM model is fitted per subject and a hierarchical PEB model with random effects is applied over the estimated connectivity parameters. Inference on network structure is implemented using a form of Bayesian Model Comparison and Bayesian Model Reduction whereby the ‘free energy’ of group level PEB models with one or more connectivity parameters switched off (i.e., prior distributions set to 0) is compared. The model ‘free energy’ represents a balance between the accuracy of a model and it’s complexity (14).

Subject-level predictions about the observed data consist of the combination of driving inputs, intrinsic connection activity, and bilinear modulation, which reflects the effects of experimental variables. In this case, the TMS stimulation served as both the driving input (on individual regions) and modulatory input (on connections between regions). These effects are modeled by the following equation (49):

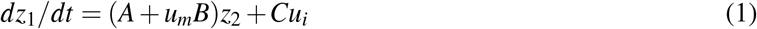

in which *dz*_1_*/dt* is the state vector per unit time for the target region, and *z*_2_ corresponds to time series data from the source region. *u_i_* indicates the direct input to the model, whereas *u_m_* indicates input from the modulatory variable onto intrinsic pathways specified by the model. Activity in the target region is therefore determined by an additive effect of the intrinsic connectivity with the source region (*Az*_2_), the bilinear variable (*u_m_Bz*_2_), corresponding to the modulatory experimental manipulation, and the effects of direct input into the model (*Cu_i_*).

In our analysis, a full DCM forward model was specified for each subject and session using the DLPFC and subgenual anterior cingulate cortex (sgACC). The DLPFC volume-of-interest (VOI) consisted of the same timeseries used as seed in the functional connectivity analysis. The sgACC VOI was defined based on the cluster of contiguous voxels within the sgACC mask that survived *p* < 0.05 uncorrected for the omnibus F-test in the functional connectivity analysis. The 1^*st*^ level DCM model was specified as follows: four experimental conditions corresponding to the same four alpha bins described in the functional connectivity analysis were included as inputs to the model; slice timing model was set to the middle slice of each volume; the default ‘one-state’ bilinear model was selected; stochastic effects were not included in the model; and time series (not cross spectral densities) were fitted because we were interested in condition specific, time-varying connectivity due to TMS and its alpha phase on stimulation onset.

For the main analysis the input matrix *U*, which models all experiment inputs across time, was mean-centered as this can increase the model evidence by enabling the connectivity parameters to stay closer to their prior expectation (of zero) during model inversion (15). We compared these results to a similar set of DCM models where the input matrix *U* was not mean-centered so that the *A* matrix could be interpreted as the effective connectivity of the unmodelled implicit baseline (i.e., time between TMS onsets) rather than as the average effective connectivity across experimental conditions (i.e., all TMS onsets). The models’ *A*, *B* and *C* matrices were fully specified for each of the four experiment conditions (alpha phase bins). In other words, both intrinsic and extrinsic connections during baseline and experimental conditions were modeled. This full model corresponds to the model formed by combining across the 1st level of each the three factors shown in Figure S.1A in SI.

The full DCM model for each subject and session was fit using spm dcm fit.m. Models for sessions that explained less than 0.05 percent of the variability in the data (assessed using spm dcm fmri check.m) were excluded from further analysis (4 and 7 models were excluded from the **HV** and **MDD** datasets, respectively). The remaining DCMs were averaged across all sessions within each subject using spm dcm average.m. These subject level models were then used to specify and estimate a group level PEB model on the B and C parameters using spm dcm peb.m in the **HV** and **MDD** datasets separately. The estimated group PEB models were then compared against reduced models where particular combinations of parameters (relating to particular connections) were switched off using spm dcm peb bmc.m. Family-wise model comparison on the output of spm dcm peb bmc was then applied to enable groups of pre-defined models (Figure S.1A in SI) to be compared using spm dcm peb bmc fam.m. Effects of alpha phase on the estimated connectivity parameters were tested with repeated measures ANOVA using the *fitrm* and ranova MATLAB routines testing for separate means between alpha phase bins. The p-values for the F statistic were adjusted for non-sphericity using Huynh-Feldt correction.

#### 5.4.5 Bold activation: GAMM

##### Data preprocessing

A 5mm FWHM filter was used to smooth fMRI data. Nuisance regressors included motion (6 motion parameters) and large motion artifact. Signal from white matter and csf was not removed as it has been suggested they have minimal impact and may lead to signal loss compared to motion regression alone when analyzing activation in event related designs (50, 51). A whole brain GLM was performed modeling a task-based design with TMS pulse timings, and nuisance regressors were removed from the data in order to generate non-noise BOLD intensities. In each run, there were 46 TMS pulses (see Figure S.4 in SI), and the time intervals between consecutive TMS pulses were randomly determined to be either 4, 5, or 6 TRs and were uniformly distributed so that there were (45/3 = 15) of each type. Ideally, the corresponding *ϕ_α_* of each TMS pulse in each run is also uniformly distributed, but due to limited sample size in each run, a uniform distribution of *ϕ_α_* is observed only when combining across both runs and subjects (see Figure S.4 in SI).

For each TMS pulse, 6 to 8 BOLD signal points were included for the GAMM analysis. Specifically, two BOLD points corresponding to two TRs before each TMS pulse were treated as baseline points and 4 to 6 (depending on the TMS pulse interval) TRs after each TMS pulse were used as non-baseline BOLD responses (see Figure S.5 in SI). The mean TMS-evoked BOLD response was calculated for each run, and the TR of the local peak response of this averaged BOLD signal during the post-TMS (non-baseline) period was used to determine the time at which the peak BOLD response would be defined for that run. The corresponding BOLD signal at this TR (time point) represented the level of TMS-evoked activation at different alpha phases of TMS onset (more details describing the BOLD peak selection process are available in section S.3 in SI). Thus the TMS-evoked BOLD activation peak is determined based on the run-averaged peak in the post-TMS period for each trial. This peak value was then normalized using eq (2):

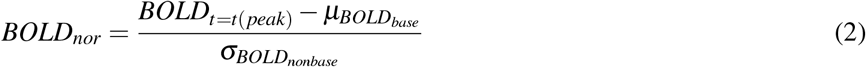

Where *BOLD_nor_* is the normalized peak BOLD signal; *BOLD*_*t*=*t*(*peak*)_ is the BOLD signal at the peak TR determined from the run-averaged TMS-evoked BOLD response; *μ_BOLD_base__base* is the arithmetic mean of two baseline BOLD signal time points before the TMS pulse; *σ_BOLD_nonbase__* is the standard deviation of the BOLD signal after the current TMS pulse but before the next TMS pulse. The normalized value *BOLD_nor_* was then used as the dependent variable in the generalized additive mixed models (GAMMs).

##### Data Structure

By merging the data from both MRI and EEG modalities, we can examine alpha phase in relation to TMS-evoked BOLD signals. For each subject, run, and region of interest (i.e. dorsal, rostral and subgenual ACC), each TMS pulse has a corresponding normalized BOLD signal and alpha phase (as calculated by the machine learning approach mentioned previously). For instance, the data structure of one subject is shown in Table 2.

**Table 2:** Dataset Structure of Example Subject

| Sub ID | $\phi_\alpha$ | $BOLD_{nor}$ | Pulse | Run | region |
| --- | --- | --- | --- | --- | --- |
| sub-01 | 1.2895 | 1.3941 | 1 | 1 | dorsal |
| sub-01 | -1.6038 | 0.4069 | 2 | 1 | dorsal |
| ... | ... | ... | ... | ... | ... |
| sub-01 | 2.4340 | 1.8892 | 46 | 6 | subgenual |

In general, for each subject, four to six runs with 46 TMS pulses per run were collected, which yielded 184 to 276 TMS pulses per session (4 to 6 runs). Trials (5%) corrupted by large motion artifact were censored (omitted) from subsequent analysis.

##### Generalized Additive Mixed Models (GAMMs)

(52) proposed generalized additive models (GAMs) to offer greater flexibility and sensitivity to detect an underlying trend between variables by allowing the use of smooth nonlinear functions to capture nonlinear relationships. It can be applied very similarly to linear regression by using smooth nonlinear functions (also referred to as nonparametric functions) in the model as parametric terms, while also providing the ability to combine the parametric and nonparametric terms in one model (53).

In general, the simplest GAM can be written as (54):

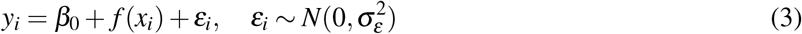

Where *y_i_* is the dependent variable at observation *i*; *β*_0_ is the intercept; *x_i_* is the predictor variable at observation *i*; *f* (*x*) is the smoothing function (e.g., *f* (*x*) = 1 + 2*x* + 3*x*^2^ + 4*x*^3^); *ε_i_* is the error term which is assumed to be independent and normally distributed. In application, smooths in GAMs are usually created with a cubic spline basis which is a preferable option compared with polynomial regressions because of their flexibility (54).

As an extension of GAMs, the generalized additive mixed models (GAMMs) adopts all features of GAMs and in addition combines the features of generalized linear mixed models (GLMMs) by adding another smooth function. The additional smooth function allows smooth terms to be treated as random effects, in addition to fixed effects (55). Therefore, we can modify GAM by adding a random effect term and write a basic GAMM as (56):

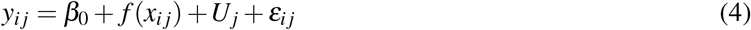

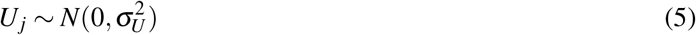

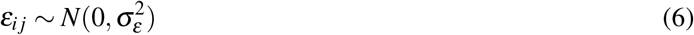

Where *y_ij_* is the dependent variable for individual *j* at observation *i*; *f* (*x_ij_*) is the smooth function of predictor variable *x* for individual *j* at observation *i*; *U_j_* represents the random effect of individual *j* which is assumed to be normally distributed; *ε_ij_* is the error term which is assumed to be independent and normally distributed.

The output of the GAMM is computed and visualized with the R Package mgcv and mgcViz (55, 57) in R studio (version 4.0.6). Based on the GAMM designed below (see eq.(8)), the smooth function of the alpha phase is specified as part of the fixed effects model formula, and intercept terms are included to provide more flexibility for the model to fit overall intercept differences between the factor levels (dACC, rACC or sgACC) and avoid artifacts due to the centering of smooths with factors. Subject differences are treated as random effects and modeled as random smoothing functions which include both random intercepts and random slope effects. Thin plate spline smooth was used for the smooth term, because in general thin plate spline is the optimal smoother of any given basis dimension/rank (58). Conventionally, thin plate regression splines do not have ‘knots’ which determines the upper bound of the number of underlying base functions being used to build up the smooth curve, so the knots of the model are not reported here (55).

In our analysis, we are interested in investigating whether the phase of prefrontal cortical alpha oscillations prior to TMS stimulation of dorsolateral prefrontal cortex (DLPFC) is correlated with neural activation within different target engagement areas (dACC, rACC and sgACC). The GAMM is established as below:

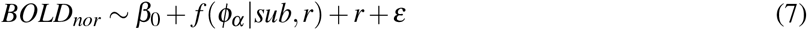

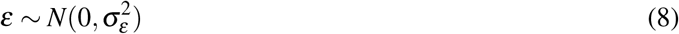

where *BOLD_nor_* represents the normalized BOLD signal; *β*_0_ is the intercept term; *f* (*ϕ_α_* |*sub, r*) is the smooth function of *ϕ_α_* (alpha phase) that has *r* (region factor: dACC, rACC, or sACC) included to model differences between target engagement areas and random effect *sub* is modeled by applying random smooths to adjust the trend of the predictor variable; *ε* is the error term that assumed to be independent and normally distributed.

## 6 ACKNOWLEDGMENTS

This work was funded by the National Institute of Mental Health (MH106775) and a Vannevar Bush Faculty Fellowship from the US Department of Defense (N00014-20-1-2027). We would like to thank Daniel Cook for his help with initial data collection with closed-loop EEG-rTMS. We would like to thank Michael Milici for his help with building the safety circuit box and ActiChamp testing. We would like to thank DeeAnn Guo for her help with ActiChamp testing and initial EEG data collection.

