## Supplement for "Functional and effective connectivity between dorsolateral prefrontal and subgenual anterior cingulate cortex depends on the timing of transcranial magnetic stimulation relative to the phase of prefrontal alpha EEG"

[1]Department of Psychiatry, Columbia University Irving Medical Center, New York, NY 10027, USA
[2]Department of Biomedical Engineering, Columbia University, New York, NY 10027, USA
[3]Department of Orthopedic Surgery, Columbia University Irving Medical Center, New York, NY 10032, USA
[4]Department of Radiology and Radiological Science, Medical University of South Carolina, Charleston, SC 29425, USA
[5]Department of Neurology, University of Chicago, Chicago, IL 60637, USA
[6]Center for Biomedical Imaging, Medical University of South Carolina, Charleston, SC 29425, USA
[7]Department of Computer Science, Columbia University, New York, NY 10027, USA
[8]Department of Radiology, Columbia University Irving Medical Center, New York, NY 10032, USA
[9]Department of Psychiatry and Behavioral Sciences, Medical University of South Carolina, Charleston, SC 29425, USA
[10]Center for Healthy Minds, University of Wisconsin-Madison, Madison, WI 53705, USA
[11]Ralph H. Johnson VA Medical Center, Charleston, SC 29401, USA
[12]Department of Electrical Engineering, Columbia University, New York, NY 10027, USA
[13]Data Science Institute, Columbia University, New York, NY 10027, USA
*contributed equally

February 15, 2022

### S.1 Supplementary Results

*ROI analysis:* In both primary and replication datasets, a main effect of TMS was observed in dACC (p<0.05 FDR corrected), but not in rACC or sgACC (data not shown). No effect of phase bin was detected in any of the ROIs in either dataset (corrected p-values>0.2). Figure S.6 below shows the FIR model time courses, averaged across subjects, for each alpha phase bin, ROI and dataset.

*Small Volume Correction:* Consistent with the ROI analysis, main effects of TMS were observed in dACC using both voxel-wise and cluster-extent correction for multiple comparisons (p<0.05 corrected). Significant clusters (but not voxels) were observed in rACC (the dorsal portion bordering the dACC mask) in the primary dataset, but not in the replication dataset. No main effects of TMS were found in sgACC in either dataset. No effects of phase bin on activation were detected in dACC, rACC or sgACC in either the primary or replication dataset.

*Parametric coefficients of GAMM:* The results of significance of parametric coefficients of GAMM for both **HV** and **MDD** groups are shown at Table S.1. According to the results, we can see that for **HV** group, there is no significant difference of maximum BOLD activation across regions (rostral-dorsal: $p = 0.3875$, subgenual-dorsal: $p = 0.2225$), while for **MDD** group, there is significant difference of maximum BOLD activation across regions (rostral-dorsal: $p = 0.0371$, rostral-dorsal: $p = 0.0274$).

| **HV** | $p$-value | **MDD** | $p$-value |
|---|---|---|---|
| (Intercept) | 0.0285 (*) | (Intercept) | 5.87e-08 (***) |
| rostral | 0.3875 | rostral | 0.0371 (*) |
| subgenual | 0.2225 | subgenual | 0.0274 (*) |

Table S.1: GAMM results in **HV** and **MDD** datasets and approximate significance of parametric terms. (**) indicates significance at the 99% confidence level; (*) indicates significance at the 95% confidence level; (.) indicates significance at the 90% confidence level.

### S.2 Supplementary Discussion

The rationale for the three ACC regions are described below.

*Dorsal anterior cingulate cortex (dACC)*: The dACC is involved in domain-general regulatory networks, cognitive control, and conflict and error detection and resolution. A recent meta-analysis found it was specifically involved in cognitive reappraisal and fear extinction, two different approaches to emotion regulation with relevance to depression [1]. Smaller dACC and rACC cingulate gray matter volumes (but not sgACC) have also been found in patients who who failed to remit following pharmacotherapy for depression remitted [2]. A previous study by our group found dACC was activated by single pulse TMS to DLPFC [3]. It has been suggested that an anti-depressant mechanism of TMS to DLPFC impacts cognitive control of emotion through modulating activity of the dACC [4].

*Rostral anterior cingulate cortex (rACC)*: The rACC, located at the genu of the corpus callosum, has been associated with treatment outcome in depression. Previous studies of regional glucose metabolism using positron emission tomography (PET) found greater rACC glucose metabolism predicted response to pharmacotherapy in depression [5–7]. Resting-state functional connectivity (rsFC) of the rACC with the salience network was also recently found to predict reductions in depressive symptoms, irrespective of treatment type [8], and rACC volume predicted response to an internet-based cognitive behavioral therapy (CBT) intervention, while SCC volume did not [9].

*Subgenual anterior cingulate cortex (sgACC)*: The sgACC plays an important role in the anti-depressant effect of TMS [10, 11]. A substantial literature suggests that reductions in activity in the sgACC are an integral part of the depression network and may predict eventual antidepressant effect [12–15]. Repetitive TMS (rTMS) likely exerts anti-depressant effects by decreasing the activity in the sgACC [16, 17]. Moreover, resting state functional connectivity (rsFC) between the DLPFC stimulation site and sgACC predicts treatment response to TMS, such that individuals with more negative DLPFC-sgACC rsFC respond better to the treatment [10].

### S.3 Definition of TMS-evoked peak BOLD signal

There are 46 TMS pulses within each run and 4 to 6 runs within each session (see Figure S.4. BOLD signal event-related averages with respect to each TMS pulse onset were created within each run. The 2 TRs before each TMS pulse onset are treated as baseline ($BOLD_{base}$) and BOLD signal between the TMS pulse onset and the next TMS pulse are treated as post-TMS activation ($BOLD_{nonbase}$, see Figure S.5). For this analysis we assume the peak BOLD signal occurs at a constant (session-specific) interval after each TMS onset. To average out trial-to-trial fluctuations due to noise, the session-specific interval from TMS onset to the peak BOLD signal for each ROI was determined based on the the post TMS onset BOLD event related average ($\mu(BOLD_{nonbase})$). For example, if the maximum value of the BOLD event-related average for run#1 is at 3 TRs after the TMS onset, then the BOLD signal value at 3 TRs post TMS-onset is used to define the TMS-evoked BOLD activation for each trial in run#1, see Figure S.5). The approach assumes the interval to the peak activation is fixed for the entire run.

### S.4 MDD Patient Sample Description

This is an interim blinded analysis of an ongoing clinical trial. All data for this randomized, double-blind, active comparator-controlled clinical trial (ClinicalTrials.gov ID: NCT03421808) was collected at the Medical University of South Carolina, SC, USA. Usable baseline fET data from 17 patients were available for inclusion in the analysis. During enrollment, all patients were randomly assigned to experiment or control intervention group before treatment. The inclusion criteria included diagnosis of unipolar MDD in a current major depressive episode, Hamilton Rating Scale for Depression (HRSD) score $\geq$ 20, age between 21 to 70, and fixed and stable antidepressant medications for 3 weeks prior to and during the trial. Patients also needed to show a moderate level of resistance to antidepressant treatment, defined as failure of one to four adequate medication trials, or intolerance to at least three trials. The most important exclusion criteria were that patients had to have a recordable alpha frequency in their scalp EEG, and to be able to undergo a 3T MRI scan safely. At the time of the analysis, no study participants were excluded based on this criterion. To ensure that baseline level of depression severity was stable at the time of study enrollment, patients were dropped from the study if they showed more than 30% improvement in the HRSD score from the time of their initial screening to the baseline assessment. A full list of inclusion and exclusion criteria can be found on ClinicalTrials.gov. (https://clinicaltrials.gov/ct2/show/NCT03421808). This study was reviewed and approved by the Institutional Review Board of Medical University South Carolina and written informed consent was obtained from all study participants prior to enrollment.

### S.5 Figures

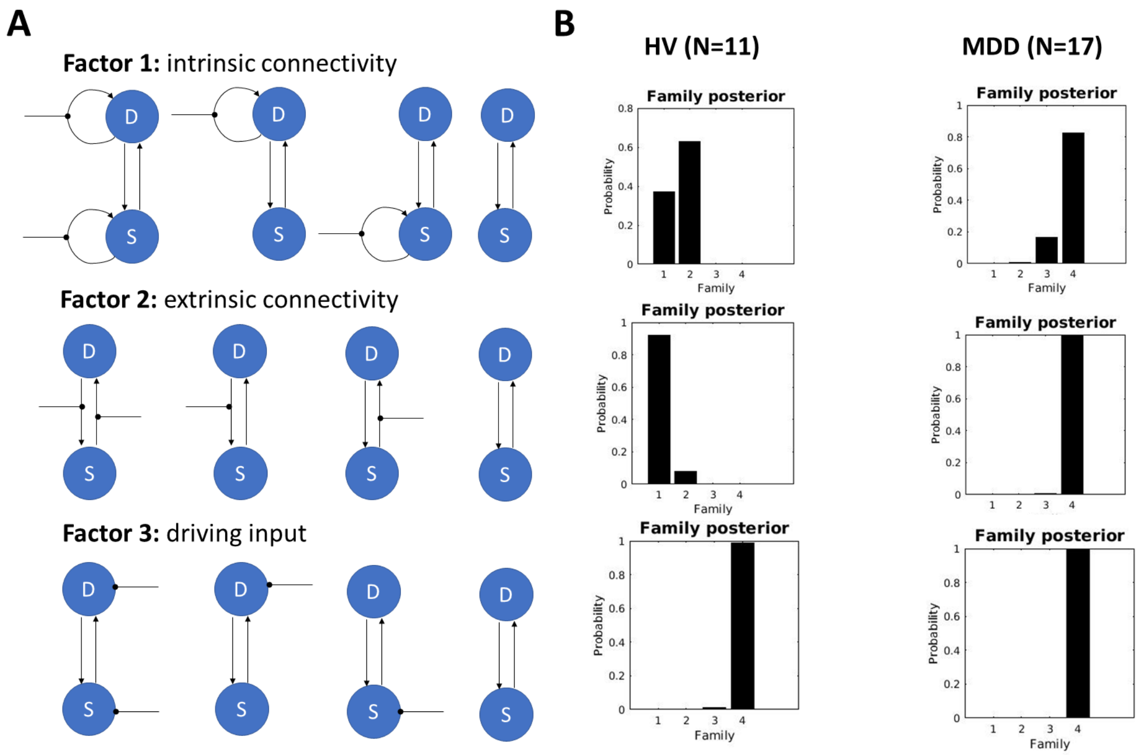


Figure S.1: Single pulse TMS to DLPFC: dynamic causal modeling with sgACC and EEG alpha phase effects. **A)** Inference on network structure was conducted using Bayesian Model Comparison and two level Parametric Empirical Bayes (PEB) for DCM. Model evidence was compared along 3 factors: TMS modulation of DLPFC and sgACC intrinsic connectivity (top), TMS modulation of extrinsic connectivity between DLPFC and sgACC (middle), and TMS as driving inputs on DLPFC and sgACC (bottom). The model space consisted of a total of 64 models, including a null model where TMS does not inform the model at all. B) Model evidence summed and plotted for each of 4 levels of the 3 factors. Factor 1 results suggest TMS modulates DLPFC intrinsic connectivity (strong evidence) as well as sgACC (moderate evidence) in HV, but not in MDD. Similarly, Factor 2 results suggest TMS modulates both top-down and bottom-up connectivity in HV, but not in MDD. Factor 3 results suggest TMS was not a driving input to either region.

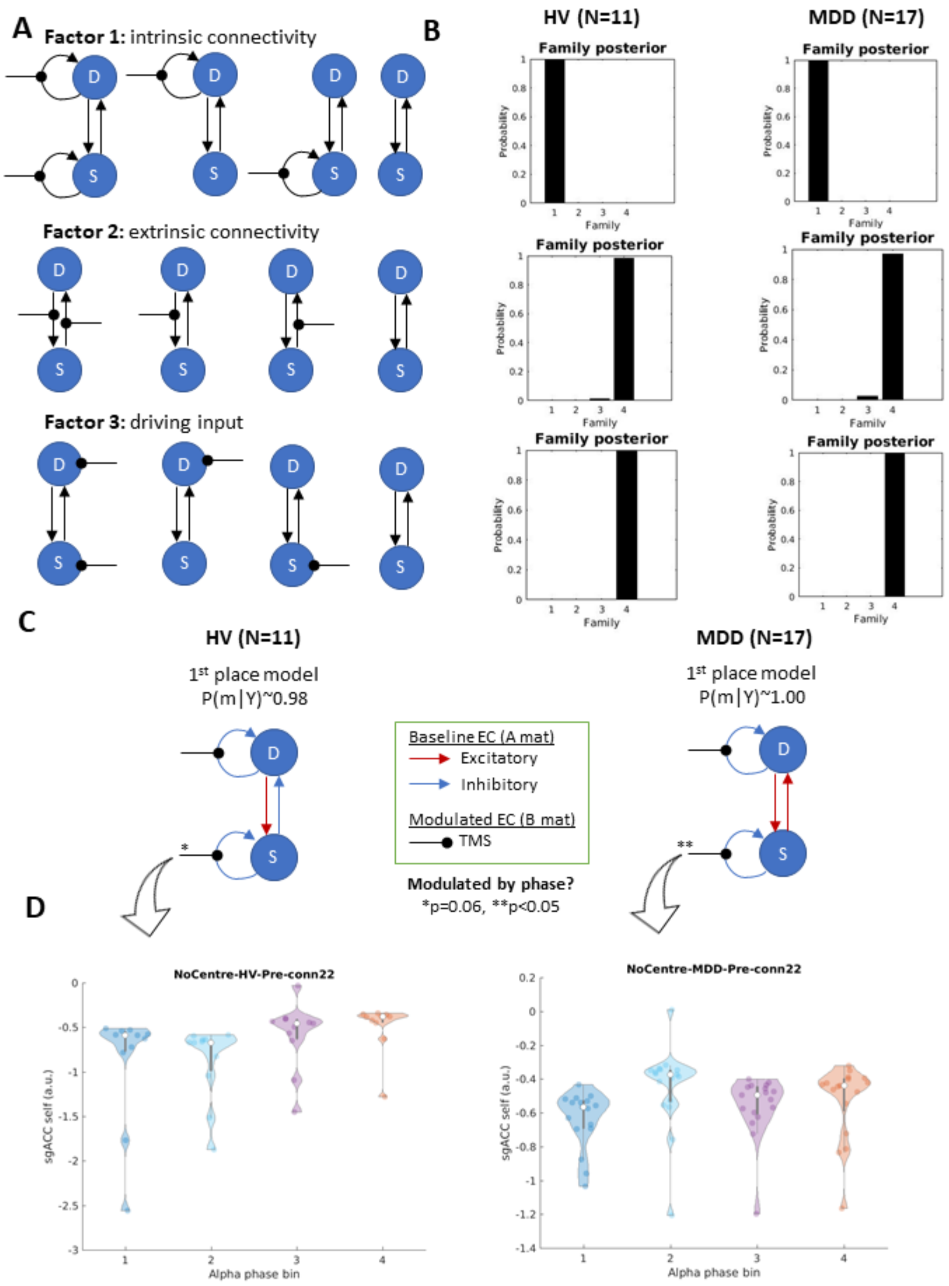


Figure S.2: Single pulse TMS to DLPFC: dynamic causal modeling with sgACC and EEG alpha phase effects with non-centered input matrix ($U$). **A)** Model space is same as shown in S.1. B) Model evidence summed and plotted for each of 4 levels of the 3 factors. Factor 1 results suggest TMS modulates DLPFC and sgACC intrinsic connectivity in HV and MDD. Factor 2 and 3 results show little to no evidence that TMS modulated top-down extrinsic connectivity in or was a driving input to either region. **C)** The top winning network structure from Bayesian Model Comparison (see methods) in HV and MDD. Results from models when not centering input matrix U suggest prefrontal alpha phase moderates TMS-evoked sgACC intrinsic connectivity (* indicates $p = 0.06$ uncorrected, ** indicates $p < 0.05$ uncorrected) **D)** Violin plots show the lowest parameter values (less inhibitory self-connection) occur during the initial rising phase of the alpha wave.

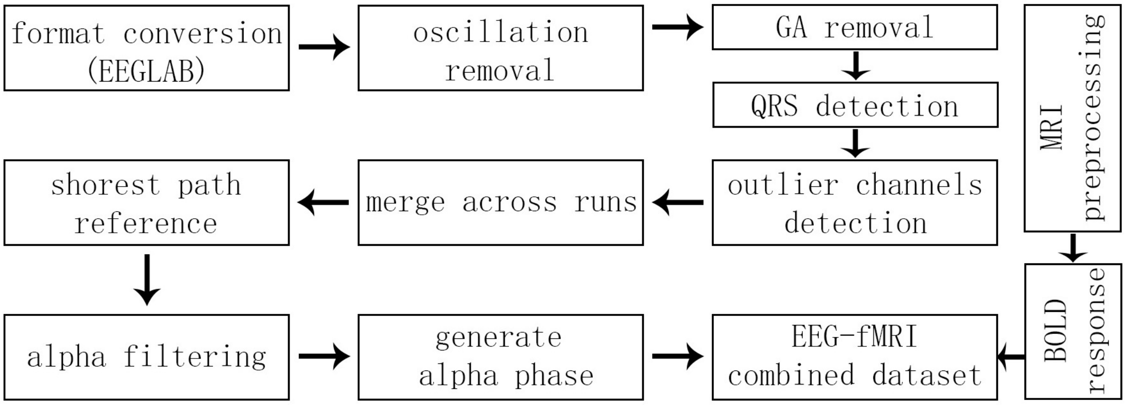


Figure S.3: Flowchart of EEG preprocessing. Each step of EEG preprocessing is listed starting from the top left. The final output (alpha phase: $\phi_\alpha$) from EEG preprocessing is combined with the BOLD activity obtained from fMRI preprocessing for subsequent statistic analysis (see Table 2 in main text for the combined dataset structure).

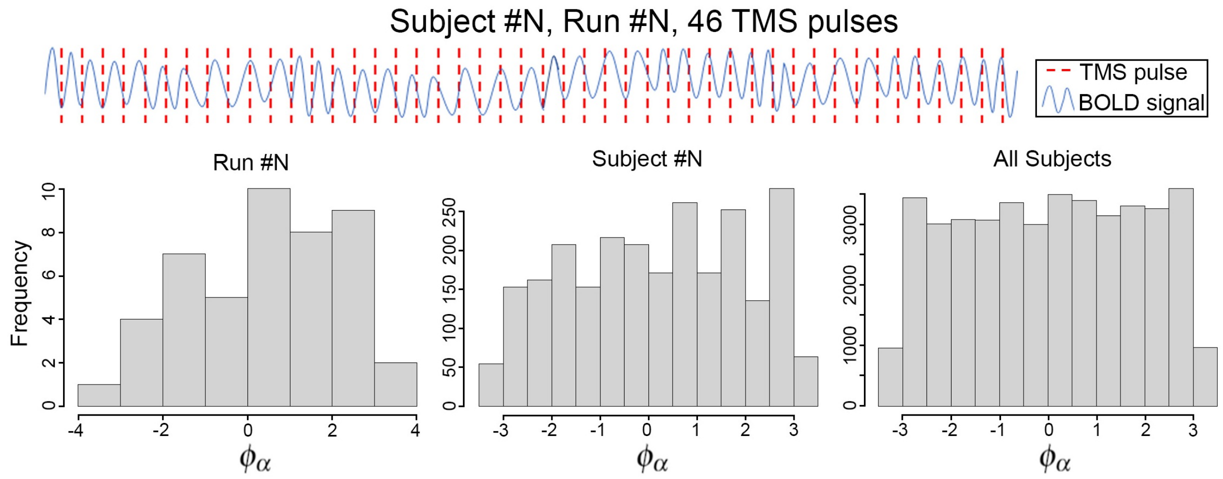


Figure S.4: Schematic of the fMRI design. Along the top is a cartoon illustration of BOLD activity from a single fMRI run with TMS pulses (dashed lines). The bottom histograms show the distribution of corresponding $\phi_\alpha$ calculated for each TMS pulse for an example run (left), an example subject/session (6 runs) (middle), and for all 17 subjects in the second dataset. In each run, there are 46 TMS pulses, and there are 4-6 runs per session. Combining data across runs and subjects yields an overall distribution of $\phi_\alpha$ that is uniformly distributed ($\phi_\alpha \sim U(0, 2\pi)$).

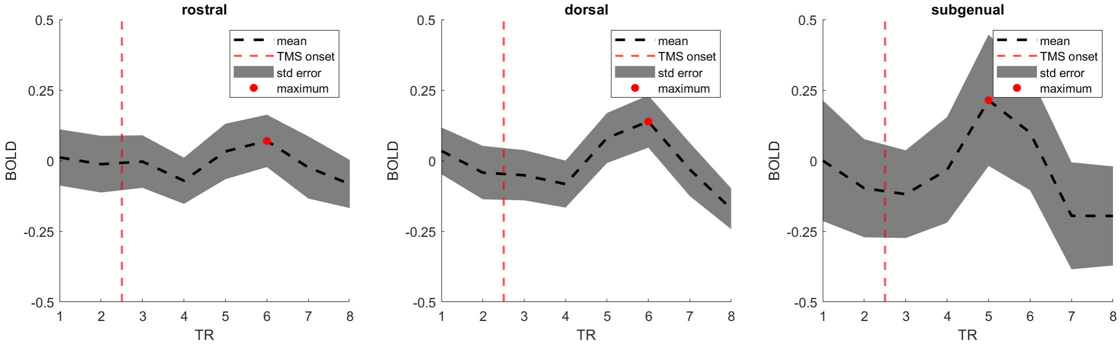


Figure S.5: TMS-evoked BOLD response definition and normalization for GAMM ROI analysis. With respect to each TMS pulse, the two scans immediately preceding the TMS onset are used as baseline ($BOLD_{base}$). All the scans following the TMS pulse and the consecutive TMS pulse are considered non-baseline/post-TMS ($BOLD_{nonbase}$) scans. The intervals between two consecutive TMS pulses are equivalently distributed among 4, 5, and 6 TRs (15 each per run). The first local peak in the BOLD response ($BOLD_{peak}$) following each TMS onset is selected to represent the TMS-evoked BOLD response. The peak is normalized using eq. (2) in the main text and is then used in GAMM ROI analysis. Sample data is preprocessed BOLD data in the 3 ACC ROIs from a single subject.

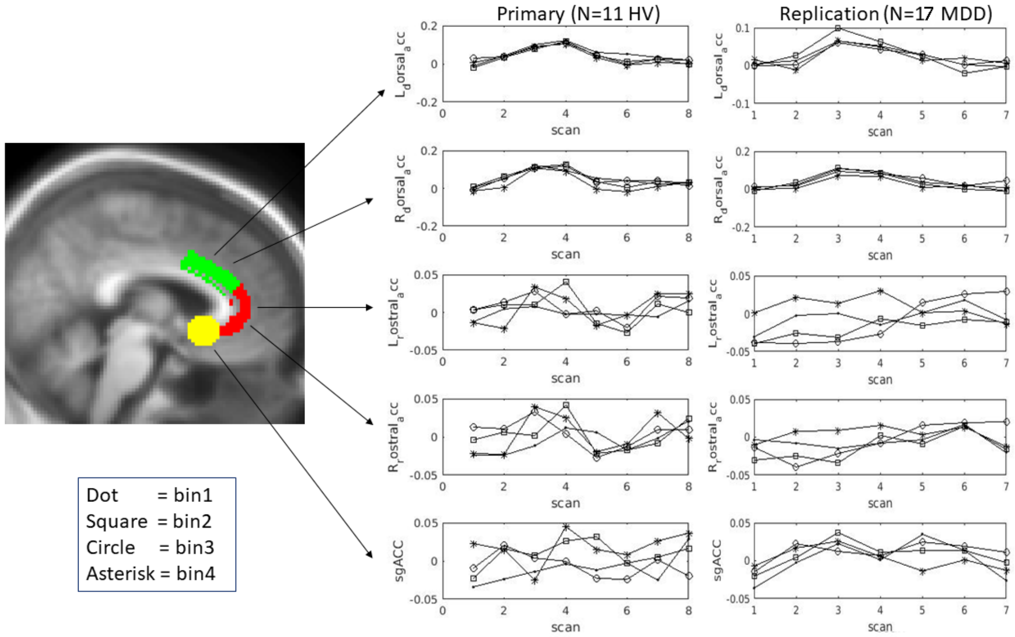


Figure S.6: Event-related BOLD signal averages across subjects for each ROI and phase bin. Average BOLD signal for each phase bins are plotted as a function of TR after TMS onset. Top row to bottom row: left dACC, right dACC, left rACC, right rACC and sgACC.
